# Inferring DNA kinkability from biased MD simulations

**DOI:** 10.1101/2025.09.08.674847

**Authors:** Arianna Fassino, Enrico Carlon, Aderik Voorspoels

## Abstract

In several biological processes, such as looping, supercoiling and DNA-protein interactions, DNA is subject to very strong deformations. While coarse-grained models often approximate DNA as a smoothly bendable polymer, experimental and theoretical studies demonstrated that mechanical stress can induce localized kinks. Here, we employ the Rigid Base Biasing of Nucleic Acids (RBB-NA) algorithm to systematically probe the properties of highly deformed DNA in all-atom simulations of a few dodecamers. A simultaneous bias in bending (roll) and twist is applied locally, to two consecutive base-pairs in the center of the dodecamers. Using umbrella sampling we construct free energy landscapes that reveal sequence-dependent effects for kink formation and quantify the energetic cost of kinking. We identify distinct features of the free energies highlighting anharmonic effects, such as asymmetries in the positive vs. negative roll. Our analysis suggests two distinct kinks characterized either by positive roll and undertwisting (twist-bend kinks) or by negative roll without excess twist (pure bend kinks). The former are frequently observed in DNA-protein structures and expected to be favored in-vivo in negatively supercoiled chromosomes. The latter have been observed in DNA simulations of minicircles and are favored in torsionally constrained DNA.

## I. INTRODUCTION

Fifty years ago, in a paper entitled “The Kinky Helix” [1], Crick and Klug proposed a geometrical model that incorporated sharp bends in DNA, or DNA kinks, to explain the folded structure of chromatin in the cell nuclei. They suggested that kinks could form in DNA without disrupting base pairing, but through the unstacking of two consecutive base pairs. According to their model, these kinks would be localized at two specific consecutive base-pairs with “bond lengths and dihedral angles assuming chemically acceptable values” [1], while the double helix would maintain is ordinary B-form at the two sides of the kink. Although sharp kinks are not essential to explain the primary features of chromatin structure – where DNA is uniformly wrapped around histone proteins – Crick and Klug’s paper sparked interest in the mechanical properties of the DNA double helix and the possibility of significant conformational changes at the base pair level.

### A. Experimental observations of kinks

Sharp kinks were indeed experimentally observed in several protein-DNA complexes [2–4]. Anomalously high bendability of bare DNA molecules, possibly associated with the presence of kinks, was reported in DNA cyclization experiments [5], atomic force microscopy (AFM) [6], DNA minicircles [7] and fluorescence resonance energy transfer (FRET) [8, 9]. More recently, X-ray and neutron small-angle scattering measurements suggested the presence of kinks in free linear DNA in solution [10]. Although some of the experimental procedures have been criticized by some authors [11–13], there is nowadays a general consensus of experimental evidence for very high bendability of short DNA sequences [14]. This bendability goes well beyond the limit expected from standard elastic models, such as the twistable wormlike chain [15].

### B. Kinks in MD Simulations

Sharp kinks of various structural conformations were also observed in all-atom simulations of small DNA minicircles [16–18] or highly bent short linear DNA [19]. These studies revealed different structural properties of kinks. In one of the earliest simulation studies, Lankas et al [16] observed two types of kinks, referred to as Type I and Type II kinks. Type I kinks closely resembled the structure proposed by Crick and Klug [1] and had intact base pairing. The Type II kinks, involved a single base pair disruption. Different types of kinks were observed also in simulations of supercoiled minicircles [17, 18], which used an updated force field, different from that used in [16]. Simulations of linear DNA biased to assume a strong bend by an applied constraint at the two ends of the molecule, found evidence of the presence of sharp kinks [19], predominantly of Type II. Bending free energies were estimated from umbrella sampling techniques [19].

### C. Models of kinkable DNA

In parallel, several groups developed and studied various analytical models that account for anomalous DNA bendability [12, 15, 20–22 To account for sharp kinks observed in AFM experiments some authors invoked a generic bending energy model[12, 15]

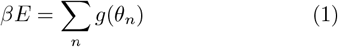

with *β* = 1*/k*_*B*_*T* and *g*(*θ*) describing the energy dependence on a local bending angle *θ*_*n*_. The total energy is then obtained by summing these local terms along the sequence. In these models the sequence is discretized in segments comprising a few helix turns, hence the sum over *n* can refer to a single base-pair step, or to a stretch of ~10 − 20 bases. Various forms for *g*(*θ*) were considered as alternative to the WLC harmonic elasticity *g*(*θ*) ∝ *θ*^2^. We mention here the examples of the linear subelastic chain *g*(*θ*) = *α*|*θ*| [6] and the hinge model

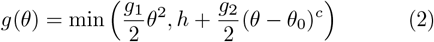

where *h* is a parameter defining the energetic cost of a kink. Two choices of exponent *c* were considered: *c* = 2 in [21] and *c* = 6 in [12]. The hinge model describes a kinked conformation as a second local energy minimum on *θ*_0_, while small bending deformations (small *θ* ≪ *θ*_0_) would be described by a harmonic term *g*_1_*θ*^2^*/*2. Yan and Marko [20] described kinks as locally disrupted base pairing causing very flexible single stranded DNA stretches, which would favor sharp DNA bending. In their model the bending energy *g*(*θ*) contains an additional “spin” variable *σ* fluctuating between the B-DNA (*σ* = 0) and locally melted, kinked (*σ* = 1) state. Although simple functions for *g*(*θ*) may be useful to provide analytical model to estimate the effect of kinks, it is desirable to gain some more insights on anharmonic effects directly from detailed MD simulations, which is the aim of this paper.

While (1) uses a single bending angle, at the base pair level there are two bending directions defined as tilt and roll. Together with twist, these describe the major deformation modes of the double helix. In the harmonic approximation the DNA energy using tilt, roll, and excess twist (*τ, ρ*, Ω) variables is given by *E* = Σ_*n*_ *ε*_*n*_, where the sum is over dinucleotide steps and with

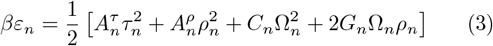

The stiffnesses 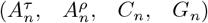 are sequence-dependent, as indicated by their *n*-dependence. The predominant off-diagonal interaction is the coupling *G*_*n*_ between roll and twist [25]. Various effects of this coupling have been discussed in the recent literature [26, 27]. A tilt deformation is much stiffer than roll 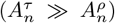, therefore large bending deformations predominantly involve roll. The presence of the twist-roll coupling suggests that a large roll should involve a twist deformation as well, consistent with observation from simulations [26, 27]. We will be seeking to infer the structure of the free energy (3) beyond the quadratic approximation.

### D. Kinks in protein-DNA complexes: the IHF

In 1998, Olson et al. [28] developed a method to extract elastic parameters of DNA deformations from high resolution protein-DNA crystal structures. They modeled DNA as a harmonic system (as (3)) and obtained sequence-dependent stiffness matrices from these structural data. Following the same line of thoughts, one can infer the kinkability of bare DNA from highly bent DNA bound to proteins. A particular interesting case, for highly bent conformations, is that of the Integration Host Factor (IHF) a DNA-binding protein found in bacteria which plays an important role in chromosomal compaction and gene regulation. IHF is a heterodimer protein consisting of two subunits which induce two sharp bends in DNA. The DNA-IHF also exhibits complex kinetics, which was studied in detail [29, 30]. Figure 1 shows (a) a structure of the IHF-DNA complex from Protein Data Bank and (b) the corresponding values of the coordinates tilt, roll and twist as calculated by the web3dna server [24] to the pdb file of the crystal structure. The two kinks are observed in TT and AA steps, with the latter intentionally nicked to facilitate crystallization of the sample [23]. The tilt, roll and twist parameters around the nicked site are not shown, as the nick breaks the B-DNA structure and such parameters do not provide useful information about the DNA conformation. The TT kink shows a strong increase in the roll (+60°) and a more moderate decrease in the twist (−10°), as expected from a positive twist-roll coupling (several simulation studies [31, 32] have shown that *G >* 0). At the kink site there is a modest (6) and the hinge model increase in tilt (+5°), suggesting that one can indeed neglect this stiff degree of freedom in analyzing DNA kinks. We will focus here on deformations involving roll and twist, as these are the most relevant parameters to describe kinks [28]. Sharp kinks in DNA-protein complexes are often characterized by a strong positive roll and under-twist, see SI for some more examples.

**FIG. 1.**
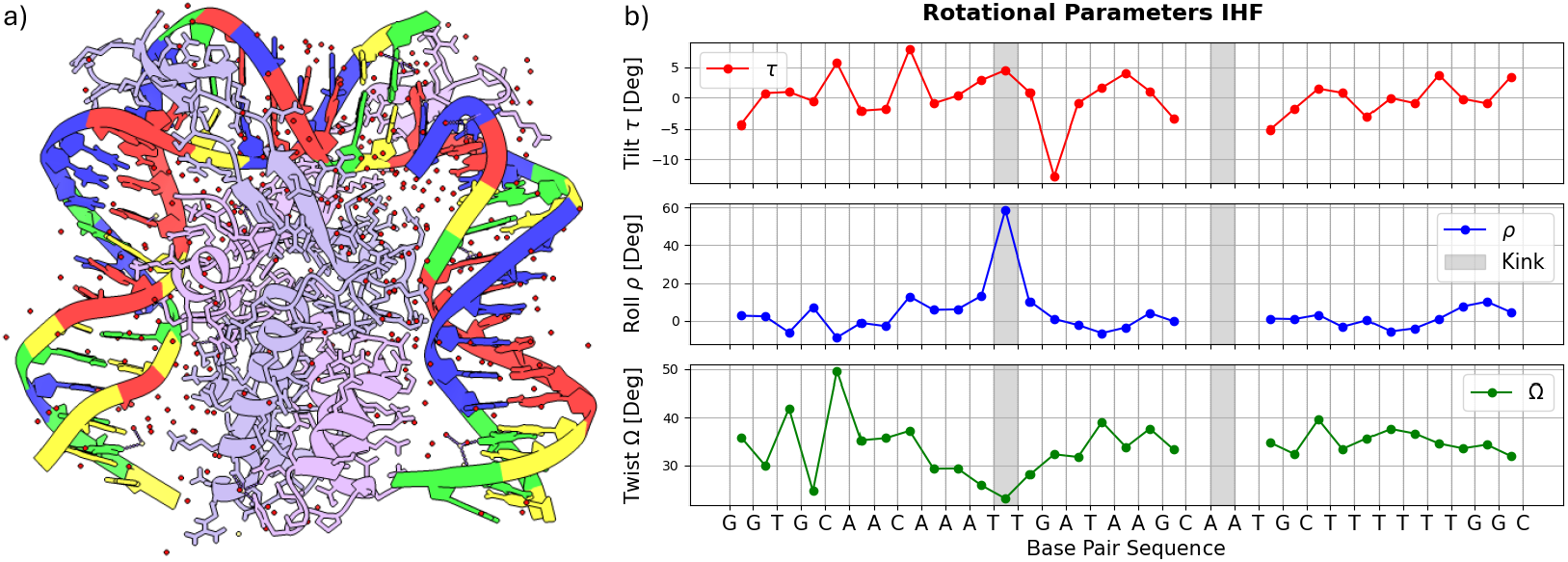
a) Crystal structure of an IHF-DNA Complex from Protein Data Bank [23] visualized via ChimeraX. The DNA forms two sharp kinks resulting in a global bend angle of ~180° over the full range of the sequence. b) Plots of Tilt (*τ*), Roll (*ρ*) and Twist (Ω) angles per base pair as obtained from the W3DNA web server [24]. The grey areas indicate the kinks location. As the DNA is nicked in the AA step (to favor crystallization) the rotational parameters are not shown. The TT-kink is characterized by a strong increase in the roll and a more moderate decrease of the twist, as expected from a positive twist-roll coupling.

### E. Kinkability from biased MD simulations

We explore here the tendency of DNA molecules to kink using all-atom MD simulations and focusing on the behavior of the roll and twist coordinates in regimes beyond the quadratic approximation (3). In unbiased MD simulations it is not possible to observe spontaneous formation of kinks, due to limitations in the timescale currently reachable in all-atom simulations. Prior MD simulations explored kinkability in DNA minicircles [16–18] which are already bent in their initial conformation. Alternatively, strong bending deformations were induced by biasing the two ends of a 15-bp linear DNA molecule [19]. In both setups kinks were seen to form in more deformable sites along the sequence and sometimes are not localized, as curvature or overand undertwisting involve several consecutive base pairs. Other MD studies [30], showed more localized kinks by simulating the entire IHF-DNA complex. This approach is crucial to capture the interplay between protein-binding mechanisms and DNA-kinkability, but cannot disentangle the two.

Here we use RBB-NA [33], a recently developed algorithm which can bias any of the 12 rigid base coordinates (or any combination thereof) at a local scale: either at a specific site or at mutiple sites. By biasing both roll and twist on a specific DNA site we infer the tendency of bare DNA to take on highly deformed conformations via the calculation of two dimensional free energy landscapes as a function of local roll and twist coordinate. The results show that kinkability is strongly sequence dependent with different sequences showing quite differ-ent behaviors. Some sequences are highly deformable and can accommodate strong and positive (roll) angles with a modest undertwist. This happens specifically for the IHF-bound DNA sequence at the kink site showing an inflection (although not a metastable local minimum) in the free energy landscape corresponding to values close to those observed in crystal structures as shown in Fig. 1(b). Other sequences however are more stiff, with a free energy increasing much more rapidly as a function of roll and twist angles. In addition we find a strongly asymmetric behavior of DNA at positive and negative rolls.

## II. MATERIALS AND METHODS

### A. Sequences used

Table I shows the six dodecamers analyzed in this work. A bias on roll and twist was applied between the two central nucleotides (in bold) of all sequences. Sequences 1 and 2 bind to the IHF protein with sharp kinks forming in the central TT and AA pairs. These two sequences are overlapping parts of the DNA of the IHF-DNA complex studied in Ref. [23] and shown in Fig. 1. Sequence 3 is part of Hbb-DNA complex, which has kinks structurally similar to those of the IHF, see [34] and SI. Histone-like protein from *Borrelia burgdorferi* (Hbb in short) is a protein which, like IHF, induces sharp bends on DNA with large positive roll and under-twist at the two binding sites (the SI reports the rotational parameters for the DNA-Hbb complex from the crystal structure data [34]). Unlike IHF, which has a high affinity for specific consensus sequences, Hbb binds DNA non-specifically, preferring binding to A-T rich regions, which are generally more flexible [34]. Sequence 4 is the Drew-Dickinson Dodecamer (DDD), a well-known sequence that has been widely studied in structural DNA investigations. Sequences 5 and 6 replicates Seq. 1 and Seq. 3, respectively, but with changes in the two biased central nucleotides with CC and GC replacing the original TT and AT. We calculated for all the six sequences the free energy Δ*F* (*ρ*, Ω) as a function of the roll (*ρ*) and twist (Ω) deformations of the two central base pairs. The protocol is schematically illustrated in Fig. 2 and explained in detail in the next Sections.

**TABLE I.** List of dodecamer sequences simulated. A bias on roll and twist is applied between the central nucleotides shown in bold to obtain free energies via umbrella sampling. Sequences 1 and 2 are IHF-binding sequences centered around the two kinks locations, see Fig. 1. Sequence 3 binds to Hbb, the Histone-like protein from the bacterium *Borrelia burgdorferi*. Sequence 4 is the Drew-Dickinson Dodecamer (DDD). Sequence 5 is as Seq. 1 with a CC pair replacing the central TT pair. Sequence 6 is mutated from the Hbb-binding sequence (Seq. 3) muted, with GC replacing the original AT central base step, instead of the natural AT step. Sequences 1-3 form sharp kinks when bound to proteins.

| Name | Origin | Sequence |
| --- | --- | --- |
| seq1 | IHF bind. | ACAAATTGATAA |
| seq2 | IHF bind. | TAAGCAATGCTT |
| seq3 | Hbb bind. | ATACTATATGTC |
| seq4 | DDD | CGCGAATTCGCG |
| seq5 | IHF mut. | ACAAACCGATAA |
| seq6 | Hbb mut. | ATACTGCATGTC |

**FIG. 2.**
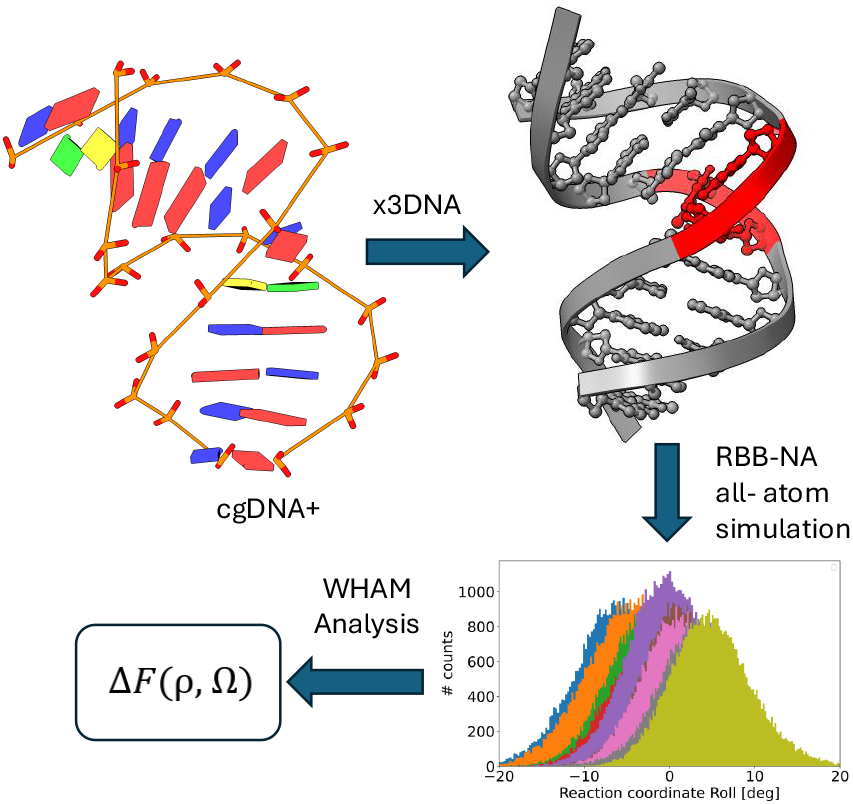
A schematic representation of the different steps followed for the calculation of the free energy Δ*F* (*ρ*, Ω). (1) The coarse-grained cgDNA+ model is biased so to produce a twisted and bemt conformation. (2) All-atom structure is then obtained by the x3DNA software. (3) Such conformations are further sampled with the RBB-NA algorithm which keeps the biases on the central nucleotides. (4) The WHAM analysis collects the data from the different biases to reconstruct the unbiased free energy.

### B. Biasing roll and twist in cgDNA+

cgDNA+ is a coarse-grained model developed to simulate sequence-dependent mechanical and structural properties of double-stranded DNA [35–38]. The standard cgDNA model [35] parametrizes the DNA conformations using the canonical twelve coordinates defined by the Tsukuba convention. Six of these are intra base pair coordinates (buckle, propeller, opening, shear, stretch, stagger) and six are inter base pair coordinates (tilt, roll, twist, shift, slide, rise). In cgDNA+, the upgraded version [36, 37], the phosphate groups are modeled ex-plicitly, which leads to 24 coordinates per bp. cgDNA and cgDNA+ sample the conformational spaces using a Monte Carlo algorithm using a quadratic free energy model for the coarse-grained coordinates, similar to (3) but with coupling extending to further neighbors (as *τ*_*n*_*τ*_*n*+*k*_ with *n >* 0). The cgDNA+ shows an excellent agreement with all-atom data, capturing some next-neighbors correlations which are missed by the earlier cgDNA model [39]. We biased the cgDNA+ model by adding the following potential

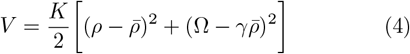

with *ρ* the roll and Ω the excess twist, as in Eq. (3). This constraint is applied to the central nucleotide pairs of the sequences of Table I and enforces roll and excess twist to assume values close to 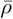 and 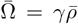, respectively. A value of *K* = 1000 kJ/mol was used. Equally spaced values (0.1 rad) of roll ranging in the interval 1.6 rad 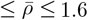 rad with *γ* = −0.1, −0.3, −0.5, −0.7, −0.9 were used. In degrees this corresponds to 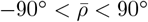. With the above given parameters we generated biased potentials for umbrella sampling [40, 41]. The two dimensional free energy landscape of the cgDNA+ model was computed using the the Weighted Histogram Analysis Method, described below, to merge data from different runs. The configurations obtained from biasing cgDNA+ were also used as input for all-atom simulations and transformed into pdb files for all-atom simulations using the x3DNA software [24, 42].

### C. All-atom system preparation

The standard Molecular Dynamics procedure includes the preliminary steps here summarized. All work is done using version 2022.5 of Gromacs, and version 2.8.3 of PLUMED. The topology file was created using Amber99 parmbsc1 force field [43, 44]. The DNA strands were placed into a dodecahedral box, leaving 2.0 nm of extra space on both sides of the molecule, with periodic boundary conditions and water model TIP-3P [45], non bonded interactions are regulated with cut-off of 1 nm and for electrostatics the PME model is used [46]. The system is solvated in a 150 mM NaCl solution after which the overall charge was neutralized. The following passages were performed using a time step Δ*t* = 2 fs with Leapfrog integrator and employing LINCS constraints. Those steps comprehend an energy minimization with a tolerance of 1000 KJ/mol, equilibration in the NVT ensemble for 100 ps, maintaining a temperature of 300 K through a velocity rescaling thermostat, and lastly another 100 ps of equilibration in the NPT ensemble, with the temperature held constant at 300 K, and pressure fixed at 1.0 bar using a Parrinello-Rahman barostat.

### D. Biasing all-atom simulations: the RBB-NA algorithm

The Rigid Base Biasing for Nucleic Acids (RBB-NA) algorithm is designed to enhance molecular dynamics (MD) simulations of nucleic acids, such as DNA and RNA [33]. It allows one to impose specific structural deformations by biasing the rigid base parameters, such as (4). While biasing cgDNA+ is a relatively simple task, RBB-NA has to turn the bias of Eq. (4), which is on the coarse-grained coordinates, into appropriate forces acting on atoms. These biasing forces on atoms need to be imposed at every MD time step. For more details on how this is done see [33]. The RBB-NA algorithm is available as a PLUMED package [47]. Similar biasing schemes were proposed in some earlier works [48–50]. Here we run the RBB-NA under the same bias as those imposed on cgDNA+ to obtain the input conformation (same 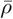 and *γ*) but with a different stiffness parameter *K*. In the RBB-NA we used *K* = 100 kJ/mol, which is 10 times smaller. A larger *K* was used in cgDNA+ as this model is considerably stiffer than the actual allatom MD at high deformations (see results), therefore one needs a stronger constraint to bias cgDNA+. A softer constraint in RBB-NA simulations allows one to sample efficiently a wider range of twist and roll values and to obtain overlapping histograms from umbrella sampling.

This in turn results in a more accurate calculation of the free energies. Figure 3 shows time traces of roll *ρ* measured during 1 ns RBB-NA simulations to illustrate the quality of relaxation under these constraints. The simulations are performed after the two 100 ps equilibration runs in the NVT and NPT ensembles, described above. On top of each plot of Fig. 3 the value of the biased roll 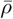 is given, while *γ* = −0.1. The roll measured in the simulation run is in absolute value smaller than the biased one, 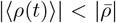. This is because the total free energy is given by the contribution of the intrinsic one (as given by Eq. (3)) plus that of the bias. The former favors the equilibrium *ρ ≈* 0. A positive/negative bias 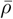 drifts the simulated average roll ⟨*ρ*(*t*)⟩ to assume positive/negative values. Most of the runs indicate that the starting point is already equilibrated, although occasionally we observe a drift, as is in the case of 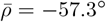 of Fig. 3. Longer runs to 5 ns however provide overlapping free energies as those from 1 ns, see Fig. S4 of the SI. In our analysis we perform multiple 32 runs for each of the five *γ*, resulting in a total of 0.16 *µ*s per sequence. The simulations with different 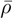 and *γ* produce overlapping histogram which are analyzed using the the Weighted Histogram Analysis Method.

**FIG. 3.**
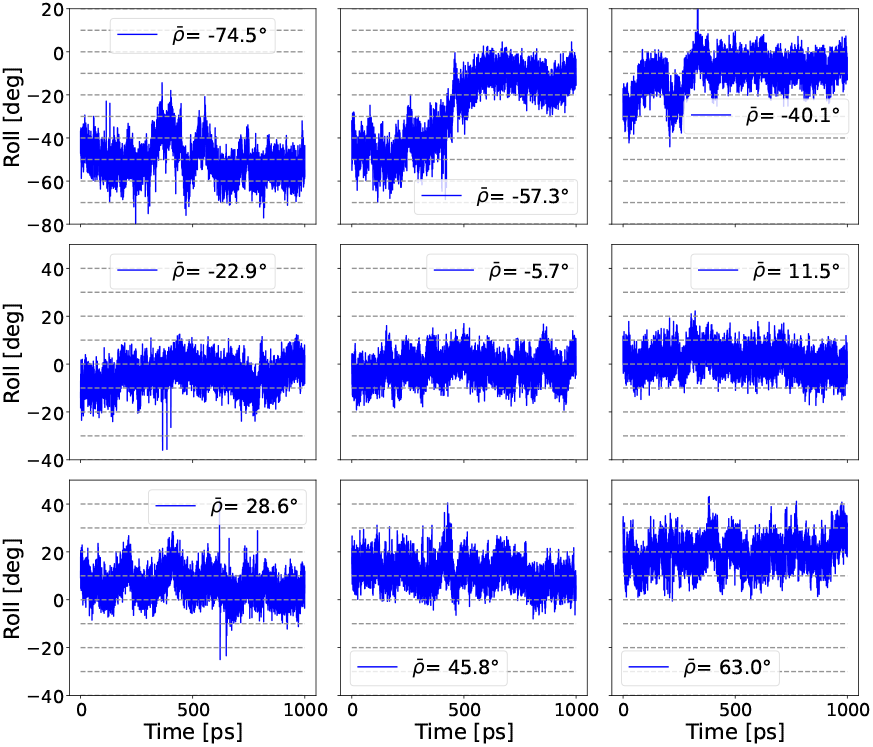
Plots of roll vs. time for 1 ns RBB-NA simulation runs for Sequence 1 with different biases 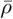, with values given in the top of each graph, and fixed *γ* = −0.1. We recall that the bias in the excess twist is 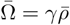. The roll *ρ*(*t*) follows the imposed bias, but note that 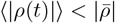. In the runs many overlapping histograms are produced from which Δ*F* (*ρ*, Ω) can be calculated. The runs indicate that they are well equilibrated with occasional slow relaxation observed at negative 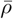, which is discussed in Results.

### E. The Weighted Histogram Analysis Method (WHAM)

The Weighted Histogram Analysis Method (WHAM) is a statistical technique through which one can merge data from multiple umbrella sampling windows to generate an estimate of the free energy profile [51, 52]. In the present work a two-dimensional free energy landscape is calculated as a function of the roll and twist coordinates. WHAM combines histograms from different biased simulations and iteratively reweights them to reconstruct the unbiased probability distribution *P* (*ρ*, Ω), from which the free energy is obtained as:

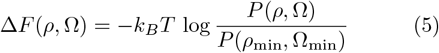

## III. RESULTS

### A. cgDNA+ free energies

Figure 4 shows a contour plot of the free energy Δ*F* (*ρ*, Ω), in *k*_*B*_*T* units, of Seq. 1 of Table I as obtained from a Monte Carlo simulation of the coarse-grained cgDNA+ model [37]. The results are given of (a) an unbiased simulation and (b) biased simulations with biasing potential given by Eq. (4). The bias values of 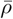 are equally spaced (with 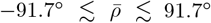 [1.6 **rad**]) and calculated along five lines of fixed slopes *γ* = −0.1, −0.3, −0.5, −0.7, −0.9. The red lines shown in Fig. 4(b) correspond to *γ* = −0.1 and *γ* = −0.9.

**FIG. 4.**
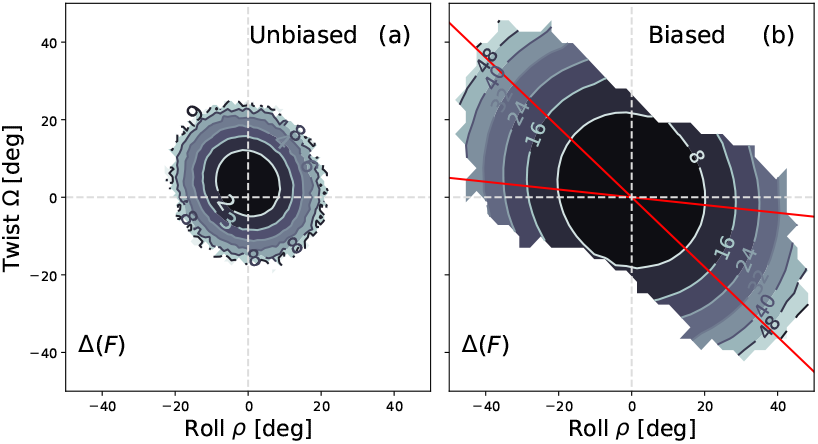
Contour plots of the free energy Δ*F* (*ρ*, Ω) for the coarse-grained cgDNA+ model for Sequence 1. (a) Unbiased simulation. (b) Biased simulation with biasing potential given by Eq. (4) for some fixed 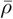 and −0.9 ≤ *γ* ≤ −0.1. The red lines indicate the smallest and largest *γ*’s. The numbers along the contour lines indicate the free energies in *k*_*B*_*T* units.

In the unbiased case (a) the range of roll and twist sampled by the simulation is limited to |*ρ*| ≲ 20° and |Ω| ≲ 20°. In the biased case (b) one can obtain a good estimate of Δ*F* for a much wider range of *ρ* and Ω. As expected, the shape of the free energy is quadratic as cgDNA+ uses quadratic terms to describe interactions between the coarse-grained degrees of freedom. The contour lines of the quadratic model are ellipses with axes which are tilted with respect of the *ρ* and Ω axes. The “tilted” landscape is due to the coupling between roll and twist discussed in the introduction (the factor *G*_*n*_ in Eq. (3)). The results of the cgDNA+ calculations are useful reference for a comparison with the all-atom free energies obtained from the RBB-NA algorithm. They are also a good test of the biasing scheme and of the performance of the WHAM, which is used to combine the results of the different simulations. The WHAM software used for cgDNA+ data analysis is the same used in the biased all-atom simulations discussed next.

### B. All-atom model free energies from RBB-NA simulations

We now turn to the results of the all-atom simulations obtained by the RBB-NA algorithm [33], as described in Materials and Methods. Contour plots of the free energies Δ*F* (*ρ*, Ω) for all six sequences are shown in Fig. 5. The data are given in units of *k*_*B*_*T*. All sequences are subject to identical bias, which is the same used to calculate the free energy of the coarse-grained cgDNA+ model of Fig. 4. The only difference is that a smaller *K* in Eq. (4) is used in the RBB-NA algorithm to allow sampling a more extended region in the *ρ*-Ω plane, see Materials and Methods.

**FIG. 5.**
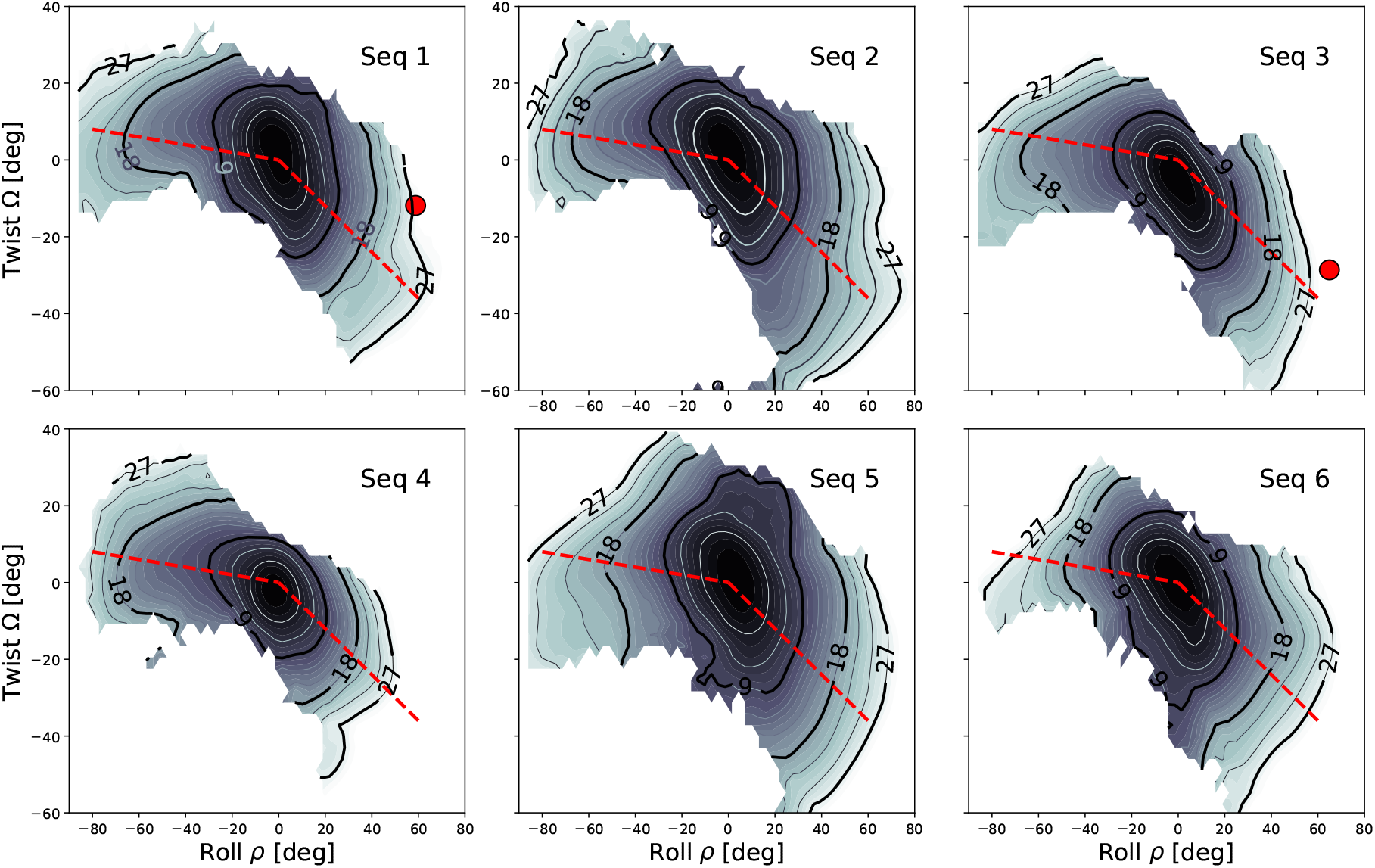
Contour plots of the free energy landscapes for the six sequences studied obtained by the RBB-NA algorithm via umbrella sampling and WHAM analysis. The free energies are given in units of *k*_*B*_*T* and the contour lines correspond to equal free energy levels, shown every 3 *k*_*B*_*T* and up to 27 *k*_*B*_*T*. The numbers printed above the lines give the correspondent free energy values. In sequence 1 and 3, the red circles indicate the values of roll and excess twist derived by the protein-bound DNA crystal structure *ρ* = 59°, Ω = −12° for Seq. 1 (IHF, pdb:1IHF) and *ρ* = 65°,Ω = −29° for Seq. 3) (Hbb, pdb: 2NP2). The red dashed lines mark the range of inclinations over which the free energy profile is depicted in 1D in Fig. 6.

#### 1. General features

We discuss first some common features of the free energies, shared by all sequences analyzed. For small roll and small excess twist the RBB-NA free energies for all sequences analyzed follow approximately the quadratic model behavior of the coarse-grained cgDNA+ model of Fig. 4(b). In RBB-NA for small deformation parameters, the landscapes appear to be tilted with respect to the *ρ* and Ω axes, due to the effect of a positive twist-roll coupling. At larger deformations, the RBB-NA free energy landscapes become asymmetric in a change *ρ* → − *ρ*, Ω → − Ω (unlike the cgDNA+ model, which uses quadratic free energies and is therefore symmetric by construction, see Fig. 4). This behavior is shared by all sequences: for positive roll the landscape remains “tilted”, but for negative roll free energies tend to become symmetric around the *y*-axis (twist). The dashed red lines are Ω = −0.1 *ρ* (*ρ <* 0) and Ω = −0.6 *ρ* (*ρ >* 0) and are drawn as guide to the eye to indicate the degree of asymmetry between the positive and negative *ρ*. In the regime of strong deformation the quadratic approximation (3) breaks down and higher terms (odd in *ρ* and Ω) are needed to describe the free energy, see Discussion. We recall that the bias is applied along lines of constant negative slope in the *ρ*-Ω plane (see Fig. 4(b)). Although the bias is symmetric in *ρ* and Ω, we note that it generates an excess twist below −40°, but hardly above 20°, showing that overtwisting DNA becomes energetically more costly than undertwisting in the high deformation regime.

#### 2. Sequence-dependent effects

We turn now to discussing sequence-specific properties of the free energies. For Sequences 1 and 3 we showed, as red circles in Fig. 5, the values of twist and roll measured at the kinks in the crystal structure of the sequences when bound to IHF (Seq. 1) and Hbb (Seq. 3), respectively. As mentioned earlier, because of the nick present in the crystal structure, these data are not available for Seq. 2 (see Fig. 1). Although our main interest is on bare DNA properties at large |*ρ*| and |Ω|, it is useful to consider kinks in protein-DNA complexes. In these complexes kinks are characterized by large positive roll and negative excess twist, corresponding to undertwisting [28]. For a more quantitative comparison between the different sequences we have plotted in Fig. 6 the free energies along the two red dashed lines in the *ρ*-Ω plane of Fig. 5. For a given |*ρ*| we note that generally the free energies are lower in the negative roll region, compared to positive roll. Curiously, Sequence 4 (DDD) has the highest free energy of all other sequences for *ρ >* 0 and the lowest of all for *ρ <* 0. There is, in general, less sequence variability for *ρ <* 0 than for *ρ >* 0. We see no signatures of local minima in our free energies, but there are in several sequences inflection points. This is close to the behavior of the hinge model (2), built on two separate free energy branches. There are more inflections in the *ρ <* 0 region than in the *ρ >* 0 region. The only two sequences with inflections in the free energy for *ρ >* 0 are those that show kinks at positive roll when bound to IHF (Sequences 1 and 2). A quantitative comparison with the coarse-grained free energies for the same sequences shows that cgDNA+ free energies are always larger than the corresponding RBB-NA free energies. In addition, as mentioned earlier, cgDNA+ free energies are quadratic in *ρ*, while apart from local “flattening” behavior at some angles, all-atom data tend to increase linearly with the bending angle. This behavior was observed in earlier simulations as well [33, 48] and is reminiscent of the linear subelastic chain behavior [6], which uses a free energy *g*(*θ*) = *α*|*θ*| in Eq. (1).

**FIG. 6.**
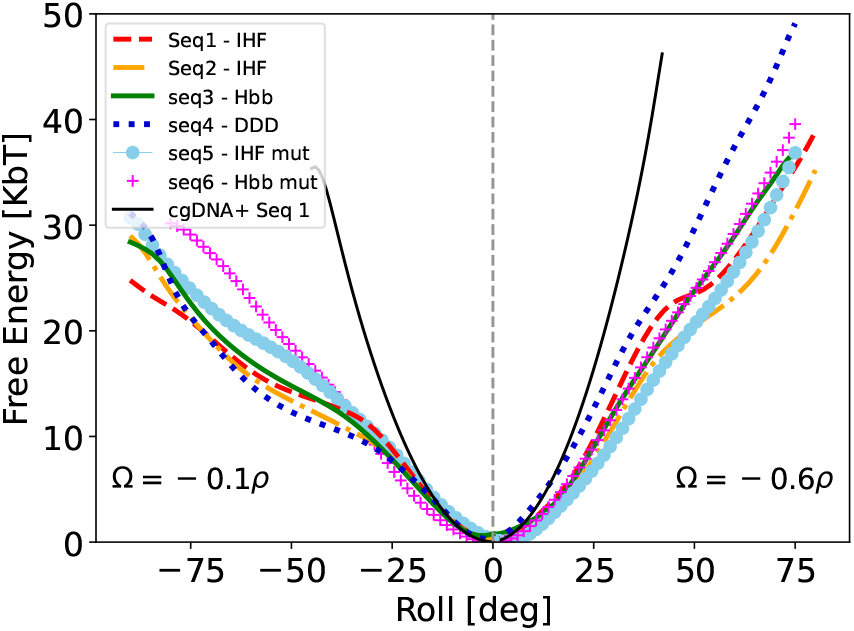
Free energies of the six sequences calculated with the RBB-NA algorithm plotted along the red dashed lines shown in Fig. 5 with slopes Ω = −0.1*ρ* (shown for *ρ <* 0) and Ω = −0.6*ρ* (shown for *ρ >* 0). For a description of individual behavior see text. The thin solid black line is the (quadratic) free energy from the cgDNA+ model for Seq. 1. This model overestimates the actual all-atom free energy data, showing that the DNA is much more deformable than predicted by the harmonic approximation.

### C. Base-pairing disruption

Forcing the system into extreme deformations with high twist and roll can disrupt base pairing in the DNA double helix in a RBB-NA simulation. Figure 7 shows two snapshots of biased RBB-NA simulations with a) intact base pairing and b) broken hydrogen bonding leading to base flipping. In the calculation of the free energy landscapes of Fig. 5 configurations with broken hydrogen bonds were excluded from the analysis. This was done as follows. During the simulation runs four additional intra base pair parameters Stagger, Shear, Stretch and Opening were monitored. Using cgDNA+ we computed the standard deviation of each parameter for every sequence as a reference value. At the end of the simulation run, using RBB-NA we then computed the mean of the parameters. Deviations exceeding four standard deviations (equivalent to a free energy cost of approximately 10 *k*_*B*_*T* under the harmonic approximation) were excluded from the analysis.

**FIG. 7.**
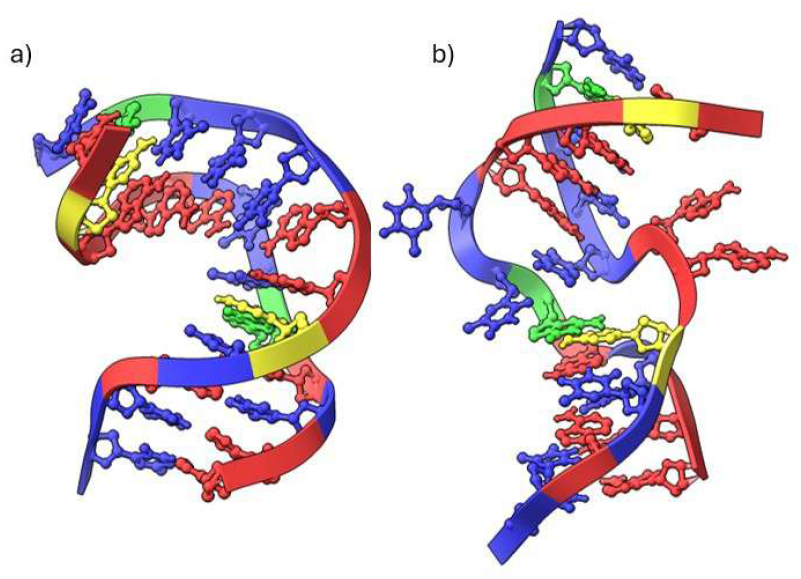
Comparison between snapshots of RBB-NA simulation runs for Sequence 1 biased to (a) 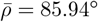 (1.5 rad), *γ* = −0.9 and (b) 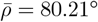 (1.4 rad), *γ* = 0.9. In the case (a) the sequence can withstand the strong bias without breaking the hydrogen bonds. In the case (b) the bias leads to hydrogen bond breakage and four bases simultaneously flipping out. Configurations of the type (b) are excluded from the WHAM analysis as our focus is to determine free energies for configurations which preserve base pairing.

We then analyzed the frequency of breakage events and found that they generally occur at extreme values of Roll, particularly in the intervals 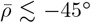 and 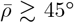 (0.8 rad). The side on which breakage occurs depends both on the sequence and on the value of *γ*. For sequence 4 and sequence 6, the majority of breakages is on the negative side whereas for the others sequences, breakages are more evenly distributed. When considering the contribution of *γ* across all sequences, we noted that for small Twist (e.g. *γ* = −0.1), most breakages are localized on the positive roll (10 in positive versus 5 in the negative). However, as the Twist increases (e.g. *γ* = −0.9) the total number of breakages rises significantly to 48, with the majority (37) laying on the negative side of roll and only 11 for positive roll. The SI describes the breakage values in more details (see Fig. S5).

### D. Beyond the harmonic approximation

The free energy data from Fig. 5 and Fig. 6 indicate that the quadratic model of Eq. (3) breaks down for roll and twist angles |*ρ*|, |Ω| ≳ 10°. To explore the nature of the anharmonic effects we analyzed the free energies using higher order powers in *ρ* and Ω via the following model

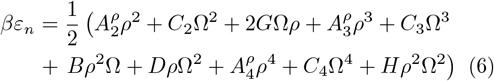

where we omit for simplicity the index *n* from *ρ*, Ω and from the stiffness parameters 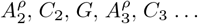 While neglecting stiff tilt degrees of freedom, the model (6) generalizes (3), by including all possible cubic terms obtained by combining *ρ* and Ω. Quartic terms were also considered, leaving out *ρ*^3^Ω and *ρ*Ω^3^. Figure 8 shows the contour plot of the free energy landscape for Seq. 4 for Δ*F ≤* 9*k*_*B*_*T* (solid lines) and the fitted data for Δ*F ≤* 7*k*_*B*_*T* (dotted lines). The harmonic model fits well free energies in the range ≲ 3*k*_*B*_*T* and we used data in this range to estimate the three quadratic parameters 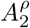, *C*_2_ and *G*. Keeping these parameters then fixed we fitted the full model (6) for the extended range of Δ*F ≤* 7*k*_*B*_*T*, with fitted stiffness parameters reported in Table II. As seen in Fig. 8 (right) the anharmonic model captures well the all-atom (RBB-NA) free energies and also the anisotropies between the positive and negative roll regions. Similar fits for other sequences are shown in SI.

**FIG. 8.**
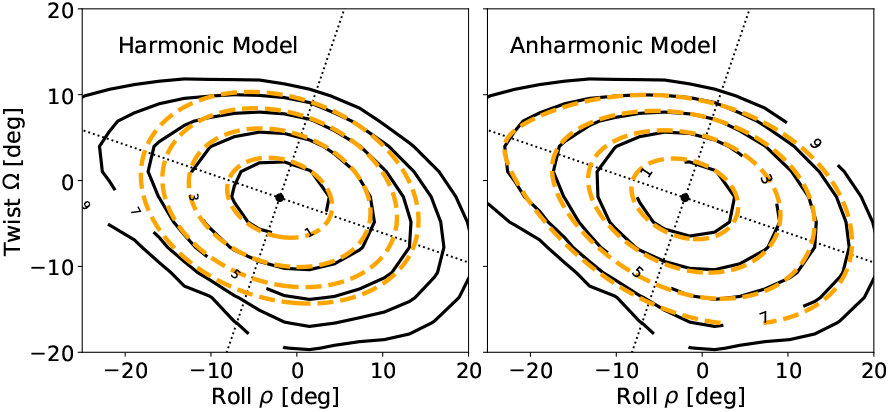
Solid black lines: Contour plots of the RBB-NA free energies for Seq. 4 (DDD). Dotted orange lines: Fitted values for the harmonic model (left) in the region Δ*F ≤* 3 *k*_*B*_*T* and for the anharmonic model (right) in the region Δ*F ≤* 7*k*_*B*_*T*. The dotted lines are the principal axes of the ellipses which are contour lines of the harmonic model.

**TABLE II.** Coefficients obtained from fitting the anaharmonic model (6) to the free energies of Seq. 4 in the range Δ*F*≤7*k*_*B*_*T*. Note: quadratic coefficients are multiplied by a factor 10^*−*2^, cubic coefficients by a factor 10^*−*4^ and quartic coefficients by a factor 10^*−*5^. The data are in dimensionless units as they are obtained by fitting angles in degrees.

| | $A_k^\rho$ | $C_k$ | $G$ | $B$ | $D$ | $H$ |
| --- | --- | --- | --- | --- | --- | --- |
| quad. (k=2) | 5.6 | 9.7 | 1.6 | - | - | - |
| cubic (k=3) | 0.2 | 10.5 | - | -2.5 | -1.6 | - |
| quartic (k=4) | -4.3 | -4.3 | - | - | - | -8.7 |

One of the remarkable features of the coefficients of Table II is 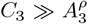 indicating the presence of a strong cubic term in the twist (~Ω^3^) as opposed to a very weak cubic roll term (~*ρ*^3^). The positive *C*_3_ suppresses over-twisting (Ω *>* 0) and favors undertwisting (Ω *<* 0) of the molecule. A large asymmetric term in twist is perhaps not surprising on view of the chirality of the DNA molecule, which is a right handed helix in its B-form. In our expansion quartic terms are negative, see Table II which lower the free energies further for high deformations. Usually, in field theory descriptions the highest-order terms must have positive prefactors which ensure that *ε*_*n*_ *→* + ∞ for large |*ρ*| and |Ω|. This stability condition is not needed in this case as the free energy (6) describes a free energy manifold of unbroken conformations which only exists for a finite range of |*ρ*| and |Ω|.

## IV. DISCUSSION

In this paper we analyzed extremely bent DNA via all-atom MD simulations. Such highly deformed conformations cannot be observed in unbiased simulations of linear DNA within the currently reachable time scales. Prior work considered simulations of DNA minicircles of 60 − 100 bp [16, 17]. These molecules are already bent in their initial conformation and this bending stress facilitates the formation of sharp kinks within relatively short time scales (~80 ns). An alternative approach was to bend a linear DNA molecule by imposing restraints at its two ends [19]. The induced bending in the latter biasing scheme is distributed along the sequence over a few base pairs and the determined free energy is associated to a global bending angle rather than to a localized deformation. Here we applied a recently developed algorithm, the RBB-NA [33], which allows one to bias any of the 12 rigid base coordinates in all-atom simulations. Although it is possible to bias multiple sites simultaneously, we have restricted our analysis here to the local biasing of the roll and twist coordinates of the central nucleotide pairs of six dodecamers.

### A. Asymmetric free energy landscapes

The main results of the simulations for the six sequences studied are given in Fig. 5, which shows the contour plot of the deformation free energies in the roll-twist plane. One of the most striking features is the asymmetric behavior in the positive/negative roll. At positive and large *ρ* the landscape remains “tilted” as is the case at small *ρ* and Ω due to twist-roll coupling. At negative and large *ρ* the tilting disappears and the landscape “aligns” along the horizontal (roll) axis, implying that bending with *ρ <* 0 does not come with a significant twist. Our results also indicate that free energies are smaller in the negative roll region, as opposed to the *ρ >* 0 region, see Fig. 6.

### B. Twist-bend kinks vs. pure-bend kinks

The landscape asymmetry suggests that there are two distinct pathways to generate sharp bending (roll) angles. The lowest free energy states for negative and large roll angles correspond to a modest associated twist. We refer to these conformations as pure-bend kinks. Conversely, the free energy landscape implies that large positive roll should have an associated negative excess twist (undertwisting). We refer to these as twist-bend kinks. The latter conformations are commonly observed in DNA-protein complexes [28], as in the example of IHF, see Fig. 1 where high positive roll is accompanied by under-twisting. While the figure shows a modest undertwisting, we note that the degree of undertwisting can also be higher than that observed for the IHF, see Figs. S1 and S2 in SI. Crystal structures of protein-DNA complexes consist of DNA fragments of ~20 nucleotides, which are torsionally unconstrained therefore could accommodate excess twist. Undertwisting is quite common in vivo, as the DNA of most organisms is negatively supercoiled, therefore it is natural to expect kinks of twist-bend type in protein-DNA complexes in living cells. The other type of kinks (which we referred to as pure-bend kinks), corresponding to large negative roll and modest or no excess twist were actually observed in simulations of DNA minicircles [16, 17]. Figure S3 in the SI shows the rotational parameters of a DNA minicircle (data from [16]) with such a strong negative roll, accompanied by a modest undertwist. We expect that this type of kinks would show up in strongly torsionally constrained DNA, as in minicircles.

### C. Strong roll/twist bias induce base-pair breakage

When the bias in the RBB-NA algorithm in the roll and twist exceeds some (sequence-dependent) threshold value, the molecule base pairs of the central dinucleotide steps get disrupted, see Fig. 7(b). However, there is a large domain of *ρ* and Ω values for which the base pairing remains intact, which comprizes large bending (roll) angles. Indeed, the free energy landscapes of Fig. 5 correspond to conformations in which the base pairs are undisrupted. The simulation data for biases which lead to base pair disruptions are not included in the free energy calculations. The kinks hypothesized by Crick and Klug [1] and also observed in DNA simulations with minicircles [16, 17] correspond to localized highly bent structure with intact base pairing. The latter simulations also generated disrupted base pairs conformations involve several (typically two or three of them), while undisrupted kinks are localized at a single base-pair step [16, 17].

### D. DNA is more bendable than the harmonic model predicts

In general we find that the free energy landscapes obtained from all-atom data are substantially lower than those extrapolated from the harmonic model (Fig. 6), indicating that the DNA is much more deformable than what predicted from the quadratic approximation. The all-atom landscapes show several inflection points which depend on the sequences analyzed and are reminiscent of the phenomenological hinge model (2), although we do not observe local “metastable” minima in Δ*F* as hypothesized in that model. For roll and twist in the range |*ρ*|, |Ω| ≤ 20°, the free energy landscapes can be fitted by a higher order model in which cubic and quartic terms in *ρ* and Ω are included, see Eq. (6) and SI. The analysis shows the presence of a strong ~Ω^3^ term favoring undertwisting, dominating over a much weaker ~*ρ*^3^ contribution.

### E. Parametrizing coarse-grained DNA models

Coarse-grained DNA models are widely used because they can reach much longer time scales and sequence lengths than all-atom simulations. Rigid base models as cgDNA [35] or analogous models [53, 54] use quadratic interaction energies and thus are expected to describe accurately DNA conformations in which the bending deformations are weak. Our analysis provide insights on the free energy landscape properties for a wide range of twist and roll values which could be used to parametrize rigid base models to go beyond quadratic approximations. Additionally it would be interesting to probe the elastic response to extreme bending deformations of other coarse-grained particle base models [55–58] and to compare them with the all-atom data reported here.

### F. Probing force fields in the high deformation regime

Our simulations use a state-of-the-art AMBER parmbsc1 force field [44], which as other DNA force fields, was parametrized for structures close to an ideal B-DNA. A systematic comparison of commonly used atomistic force fields unveiled some differences in predicting elastic properties [59]. It has been pointed out that one of the shortcomings of the current force fields for DNA simulations is the tendency to overstabilize some interaction terms in the double helix [60]. It is also believed that improved force fields will lead to a higher degree of conformational flexibility [60]. It would be interesting to test deformation free energies as those reported in Fig. 5 using alternative force fields. The RBB-NA algorithm [33] could handle these as well. It is worth noting that the value for the free energy of kinking (≃27*k*_*B*_*T*) derived form 5 is in agreement with some of the earlier all-atom MD based studies [48]. However, other authors have estimated that this value should be close to ≃15.4*k*_*B*_*T*, based on the observation that kinks compete with basepair disruption to alleviate strain [21]. This large difference may point to the fact that the kinks we find are not a stable feature of bare DNA, but are rather either transient or stabilized by proteins like IHF. Alternatively this discrepancy could indicate again that the AMBER force field used here creates too stiff DNA at high deformations.

## V. CONCLUSION

In conclusion, we investigated the behavior of highly bent DNA using the RBB-NA algorithm, through which we applied a localized bias potential to the central dinucleotide step of some selected dodecamers. The core contribution of this work lies in the detailed characterizations of the free energy landscapes. All analyzed sequences present an asymmetric behavior: DNA accommodates undertwisting through a combination of positive roll, whereas overtwisting, particularly when coupled with negative roll, results in significantly higher energetic costs, often leading to base-pair breakage. These results highlight the need for incorporating anharmonic models in DNA structural descriptions, such as the one we presented here. Furthermore, our analysis offers a pathway towards improved parametrization of coarse-grained models.

## VI. ACKNOWLEDGEMENTS

We are grateful to Davide Marenduzzo for stimulating discussions. AF acknowledges financial support by EU-funded Doctoral Network MeChaNiSM (Mechanical Characterization of Nucleic acids using Single Molecule techniques) within the Horizon Europe 2021-2027 Framework Programme Grant Agreement number 101168851. The resources and services used in this work were provided by the VSC (Flemish Supercomputer Center), funded by the Research Foundation Flanders (FWO) and the Flemish Government.

## VII. DATA AVAILABILITY

The data that support the findings of this study are available upon request. These data include: PDB files generated using cgDNA+, Scripts for running RBB-NA molecular dynamics simulations, Output files from RBBNA simulations (XTC and TPR formats), and Python scripts for WHAM analysis and figure generation.

### 1. Conflict of interest statement

None declared.

## VIII. SUPPLEMENTAL INFORMATION

### A. Kinks in protein-DNA complexes from crystal structures

It is known, as it was pointed out in early analysis of crystal structure data [**Olson 1998**], that sharp kinks in DNA-protein complexes tend to have positive roll and negative twist. Figures S1 and S2 show two additional examples (besides the IHF shown in the main text) of crystal structures of DNA complexes having each two sharp kinks for the Hbb-DNA and CAP-DNA complexes, respectively. These data show similar features as those observed in IHF: the kinks in protein-DNA structures are indeed characterized by a large positive roll and a negative excess twist. Both kinks are localized at a single basepair step, shown in grey in Fig. S1 and S2(b). The CAP-DNA complex is characterized by a smaller roll angle at the kinks *ρ* ≈ 40° as opposed to the Hbb-DNA *ρ* ≈_*°*_ 60°. In both complexes tilt varies very weakly |*τ* | *<* 10, as opposed to the strong variation in roll and twist.

**Fig. S1.**
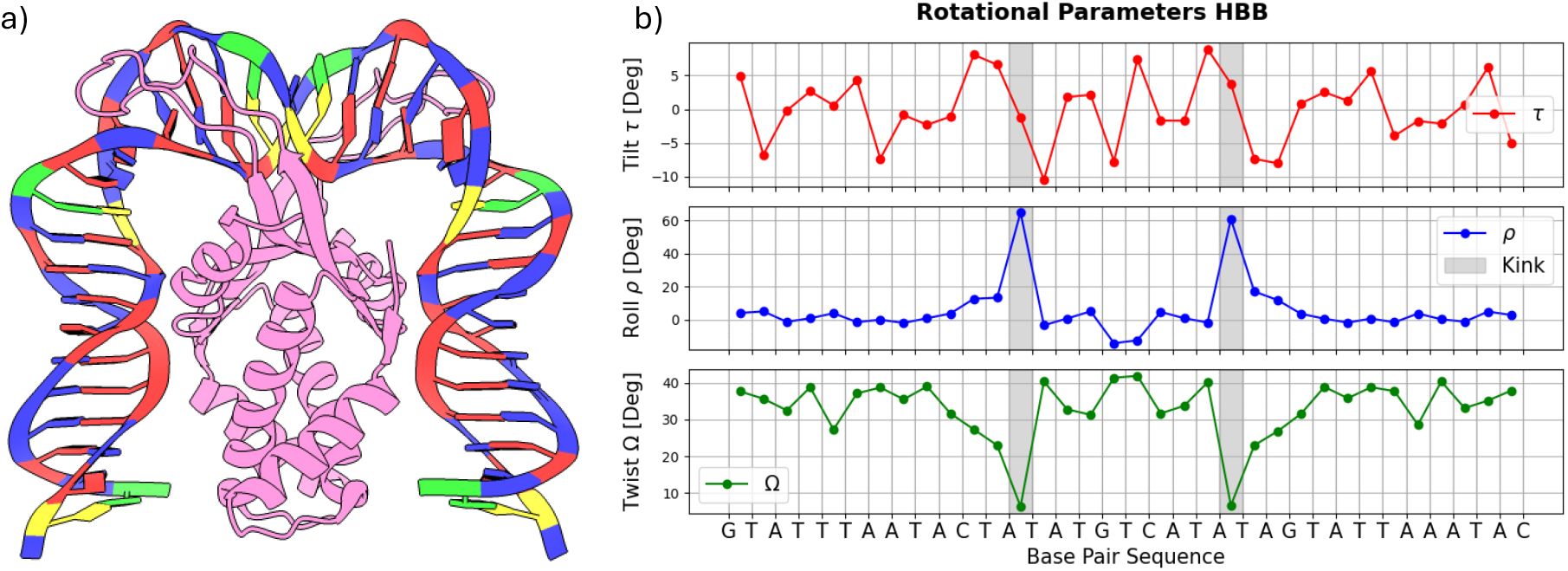
(a) Histone-like protein from Borrelia burgdorferi (Hbb in short) is a DNA binding-protein. The figure shown the three dimensional crystallographic structure of Hbb bound to DNA as from Protein Data Bank **[Mouw and Rice (2007)]**. (b) Plots of Rotational coordinate tilt(*τ*), twist (Ω), and roll (*ρ*) as obtained using W3DNA. The DNA has two sharp kinks, localized in two symmetric AT step (grey area). As for IHF (see Fig. 1) main text, the kinks are characterized by a large positive roll (*ρ* ≈ 60°) and a negative excess twist Ω ≈ −20°. Tilt varies very weakly along the sequence, with typically |*τ* | ≲ 5°.

**Fig. S2.**
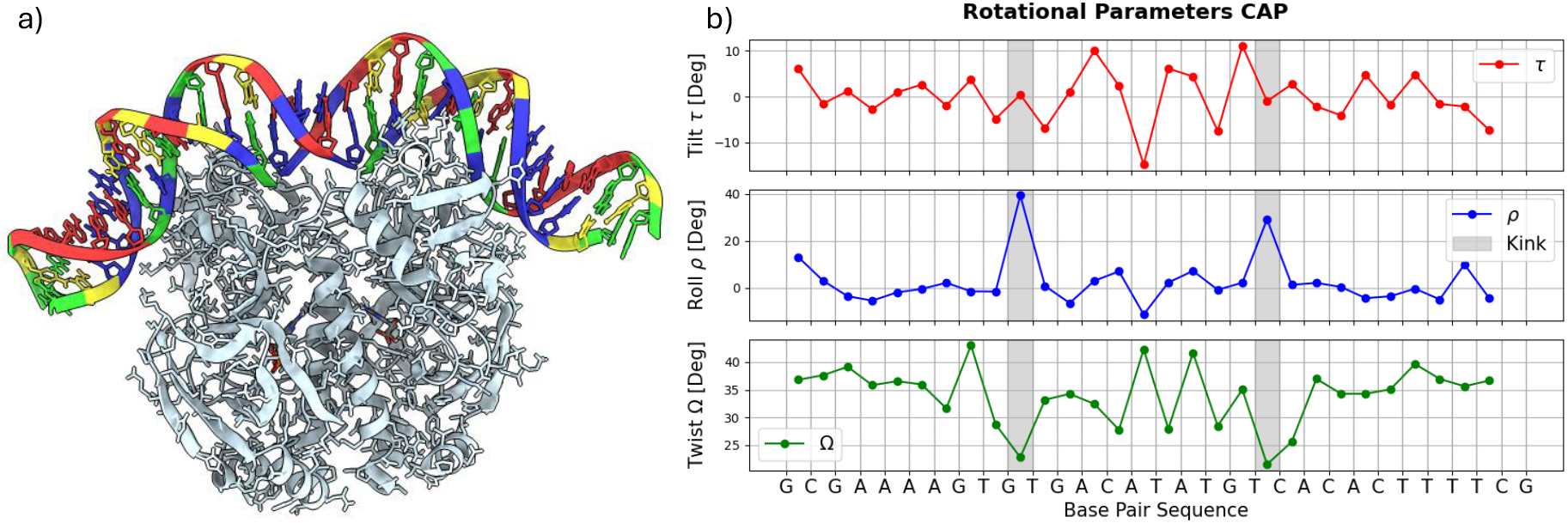
Catabolite Activator Protein (CAP) is a bacterial transcriptional activator. in E. Coli and other bacteria **[Schults (1991)]**. (a) Three dimensional crystal structure of CAP-DNA. (b) Plots of Rotational coordinate tilt(*τ*), twist (Ω), and roll (*ρ*) as obtained using W3DNA. At the two kinks these show similar features as for the IHF (Fig. 1 main text) and Hbb (Fig. S1): positive roll *ρ ≈* 40°, although not as large as the IHF and Hbb, and negative excess twist.

### B. Kinks in DNA minicircles

Simulations of DNA minicircles reported different types of kinks structures [**Lankas et al 2006, Mitchell et al 2011**]. Kinks with undisrupted base pairs were reported to have predominantly negative roll. Figure S3(a) shows a snapshot of the configuration of a minicircle of 120 base pairs after 80 ns of MD simulation. A plot (b) of the rotational parameters tilt, roll and twist shows the presence of a sharp kink with large negative roll (*ρ* ≈ −90°) and small negative excess twist. This conformation supports the report of asymmetric free energy of DNA deformations of Fig. 5 of the main paper: undertwisting is favored to overtwisting.

**Fig. S3.**
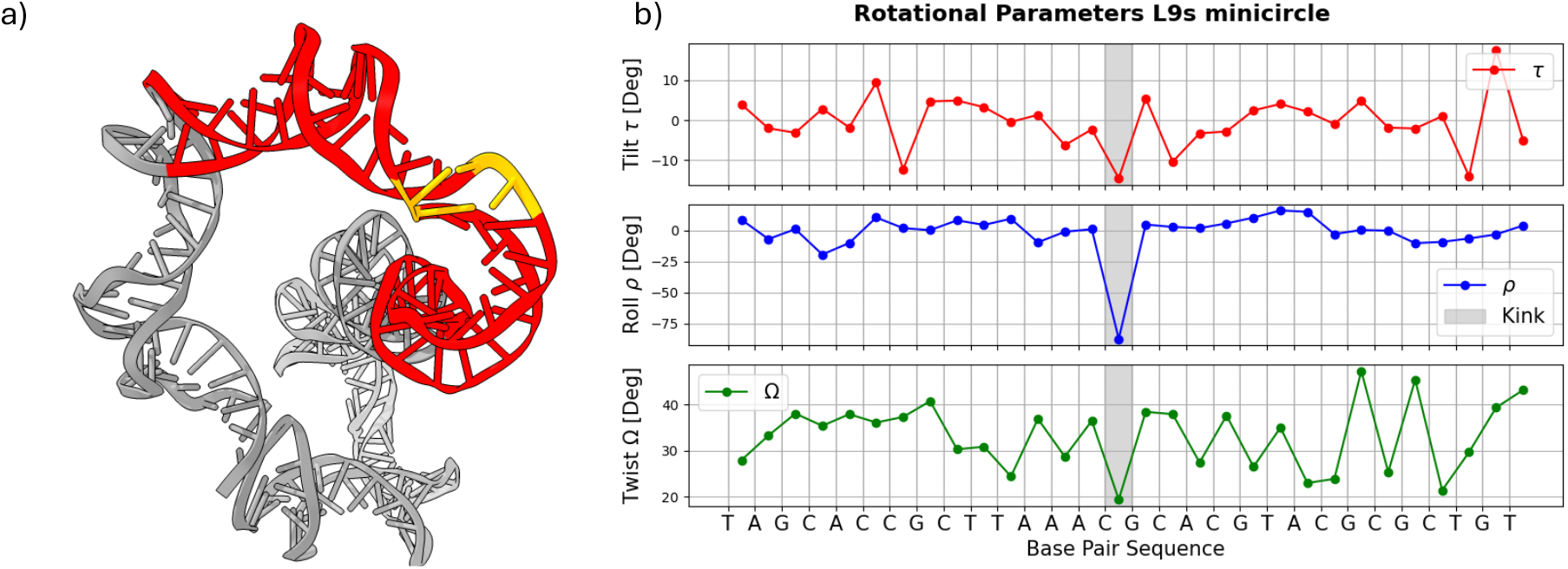
a) Three dimensional structure of Minicircle L9s as presented in the Supplementary Data in **[Lankaš et al. 2006]**. In red we show the base pairs in the vicinity of a sharp kink (yellow) for which the rotational parameters were computed. b) Plots of Rotational coordinate tilt(*τ*), twist (Ω), and roll (*ρ*) as obtained from the structure in a) using W3DNA. The type I kink is in the correspondence of CG step. We observe a negative roll (*ρ* ≃ −87°) and undertwist, corresponding to a negative excess twist (Ω ≃ −14°).

### C. RBB-NA biased runs

The Figure S4 shows some additional plots of roll (*ρ*) [deg] vs. time [ps] for various sequences and gammas. On top of each graph we printed the corresponding values of the bias applied, 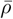. Compared to Fig. 3 of the main text here we included different sequences combined with higher *γ*, implying a stronger bias on the twist as 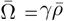. As discussed in the main text, *ρ*(*t*) fluctuates and follows the applied bias 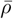, but as previously evidenced 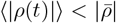. Apart from some fluctuations, the system seems to be well equilibrated after the NVT and NPT prepararion runs described in the Methodology Section of the main text. As seen also in Fig. 3 of the main, *ρ*(*t*) has stronger fluctuations for some specific values of the bias.

**Fig. S4.**
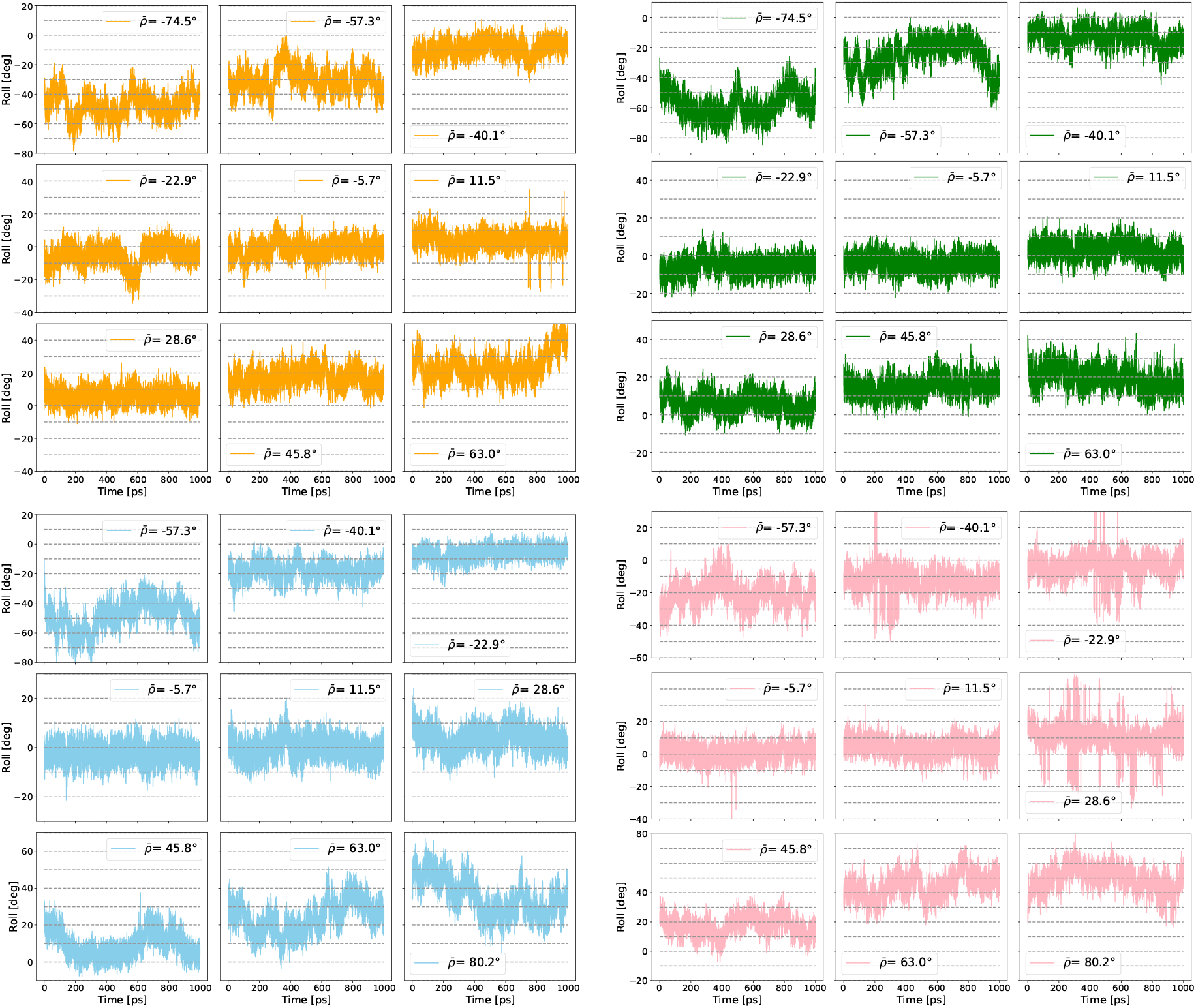
Examples of relaxation of the systems for 4 different sequences and gammas. Color code: Orange represents sequence 2 with *γ* = −0.3, Green is sequence 3 and *γ* = −0.5, Sky blue sequence 4 *γ* = −0.7, Pink sequence 5 *γ* = −0.9.

### D. Sampling free energies from longer simulation runs

RBB-NA simulation time is fixed at 1 ns for any values of applied bias 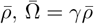. To test whether such time interval is sufficient to probe equilibrium free energies, we repeated the calculations for a 5 ns simulations for Se-quence 2 of Table 1. Figure S5 compares the free energy landscapes as obtained for 5 ns (left) and 1 ns (right). Besides a slightly wider space exploration, expected due to higher chances of exploring regions with higher |*ρ*| or |Ω| when a longer runs are performed, the two free energies are in good quantitative agreement. This shows that 1 ns runs are sufficient to sample the equilibrium free energies.

**Fig. S5.**
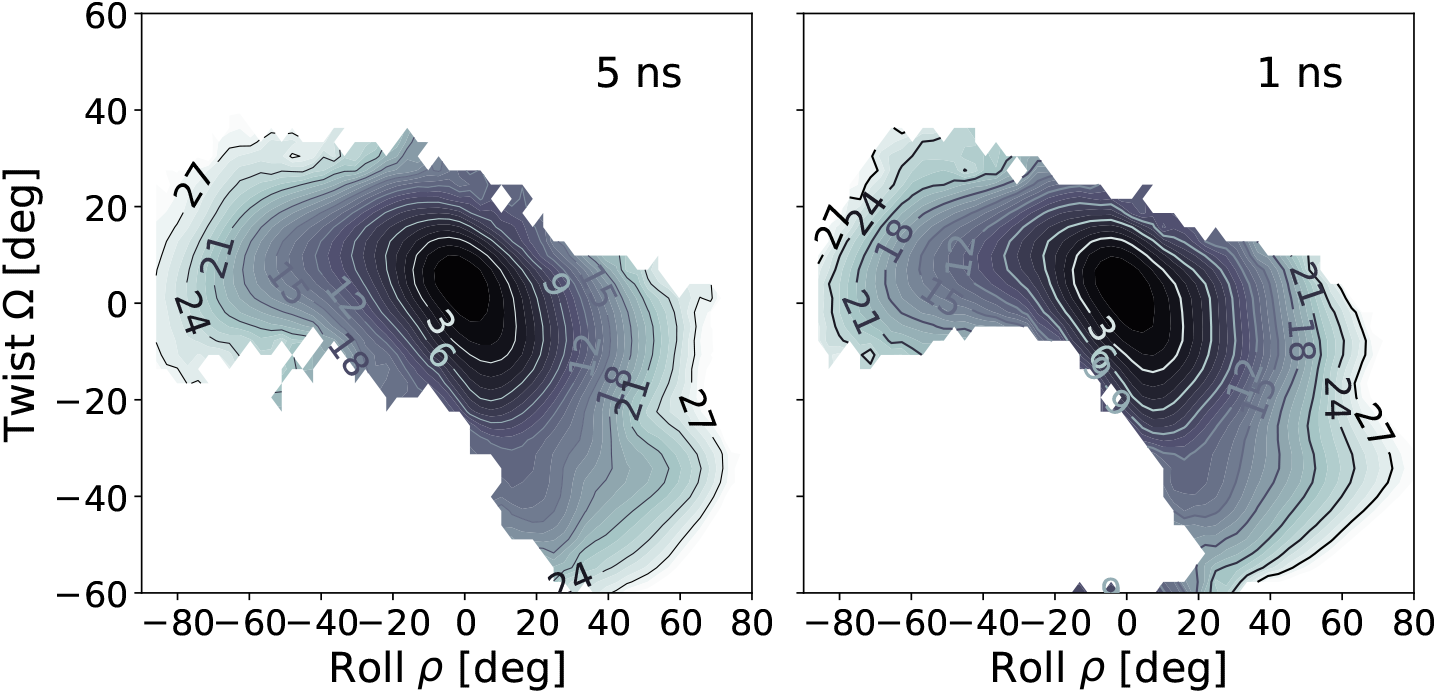
Comparison of contour plots of the free energy landscape for Sequence 2 (Kink in IHF-DNA binding) obtained by the RBB-NA algorithm with samplings of 5 ns, on the left, and 1 ns on the right. The free energies are given in units of *k*_*B*_*T* and the contour lines correspond to equal free energy levels, shown every 3 *k*_*B*_*T* and up to 27*k*_*B*_*T*. The good qualitative agreement between the two indicates that the 1 ns simulations are sufficient to sample equilibrium free energies with good accuracy.

### E. Analysis of base-pairing disruptions

In the main text, the section ‘Base-pairing disruption’ addresses this issue encountered in the MD simulation run at high biases. Forcing the system towards high values of roll and twist can lead to the breakage of basepairing. Figure S6 shows as red crosses the values of the bias of roll and twist in which we observe the disruption. Most of these crosses fall outside the range of *ρ* and Ω in which the free energy is evaluated. In a few cases they are inside the free energy domain. This happens because the criterion to flag a base-pair disruption event is based on fluctuations exceeding some threshold value (see main text). It may happen that for some given bias 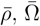 this threshold is passed and the simulation gets flagged as base-pair disruption event. Then, when the bias is slightly exceeded, it may happen that no base-pair disruptions are detected and the free energy gets evaluated. The appearance of a few isolated crosses within the free energy domain in Fig. S6 is due to some occasional false positive detection. We note that, in general, overtwisting leads to much more frequent basepairing disruptions than undertwisting. Some sequences, as Seq. 4 and Seq. 6, can tolerate a large degree of undertwisting while maintaining intact base-pairing.

**Fig. S6.**
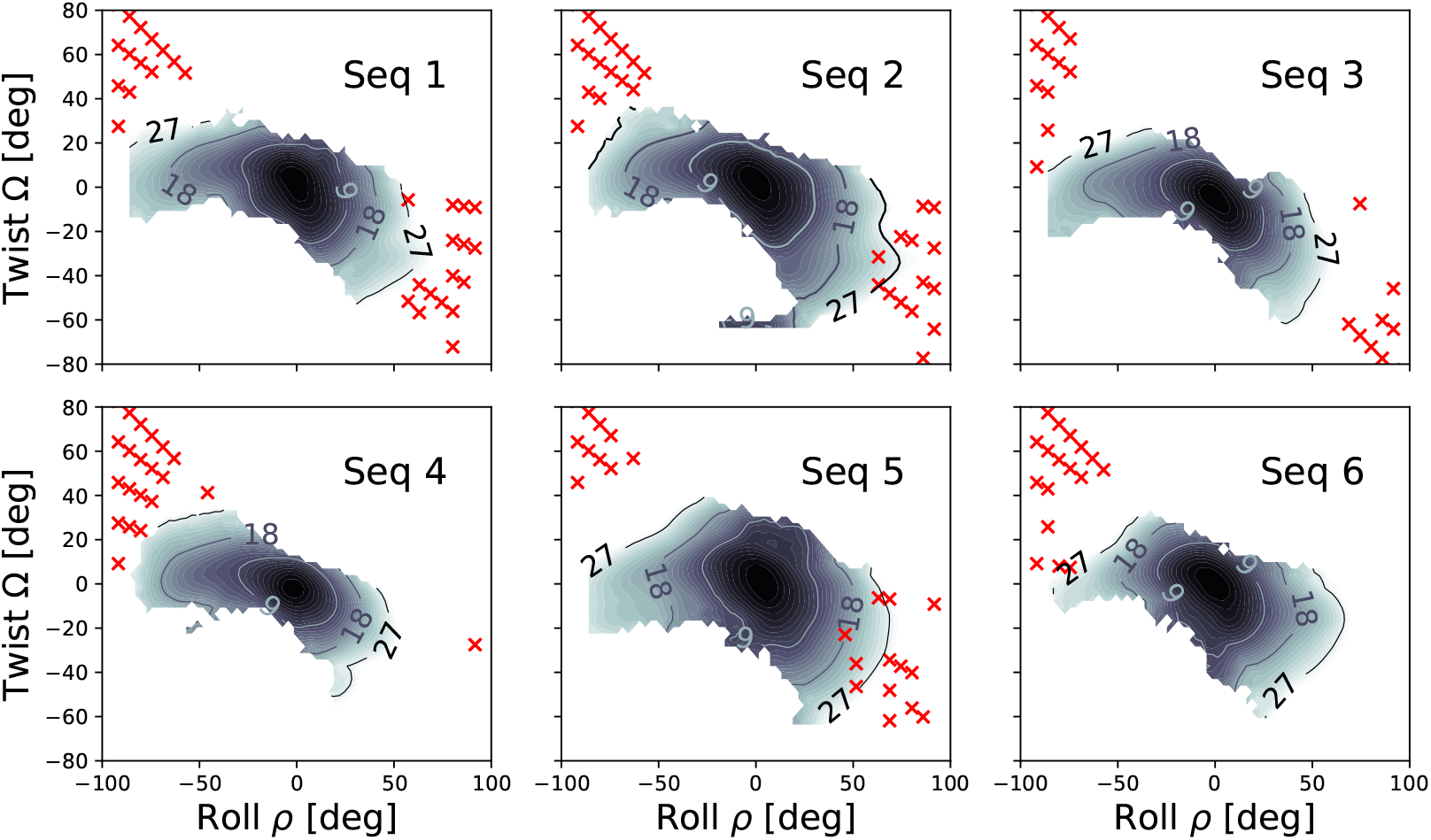
Contour plots of the free energy landscapes, for the six sequences studied, obtained by the RBB-NA algorithm via umbrella sampling and WHAM analysis. The free energies are given in units of *k*_*B*_*T* and the contour lines correspond to equal free energy levels, shown every 9 *k*_*B*_*T* and up to 27 *k*_*B*_*T*. The focus is on the configurations where base disruption was observed, indicated by the red crosses in the figure.

### F. Anharmonic Model

We discussed in the Results Section of the main text the analysis of the free energy of Sequence 4 beyond the harmonic approximation. Here, we present the analysis within the same model, described by Eq. 6 of the main text for Sequences 1, 2 and 3. The anharmonic model fits reasonable well the other sequences as well, although the quality is slightly worse for Seq. 1. Table S1 compares the fitted parameters for the four sequences. As already pointed out for Seq. 4 in the main text, one notices a large contribution from a ~ Ω^3^ as opposed to a weak 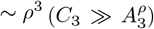. The cubic terms *B* and *D*, as well as the quartic term 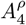 are negative in all sequences analyzed. We note that the sign of *C*_4_ and *H* however do change for the different sequences.

**Table S1.** Comparisons of fitted parameters for sequences 1,2, 3 and 4 using the anharmonic model of Eq. 6 of the main text. Quadratic coefficients 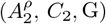 are multiplied by a factor 10^*−*2^, cubic coefficients 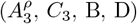 by a factor 10^*−*4^ and quartic coefficients 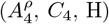 by a factor 10^*−*5^. The data are in dimensionless units as they are obtained by fitting angles in degrees.

|  | Seq 1 | Seq 2 | Seq 3 | Seq 4 |
| --- | --- | --- | --- | --- |
| $A_2^\rho$ | 4.15 | 3.49 | 4.62 | 5.61 |
| $C_2$ | 3.85 | 3.75 | 6.20 | 9.65 |
| $G$ | 1.60 | 1.92 | 3.17 | 1.57 |
| $A_3^\rho$ | 0.80 | -1.02 | 0.33 | 0.19 |
| $C_3$ | 6.28 | 10.82 | 5.37 | 10.52 |
| $B$ | -0.38 | -4.70 | -0.98 | -2.49 |
| $D$ | -4.32 | -4.12 | -4.06 | -1.60 |
| $A_4^\rho$ | -2.05 | -1.53 | -1.93 | -4.29 |
| $C_4$ | 1.90 | 3.21 | -0.52 | -4.26 |
| $H$ | 2.66 | 4.65 | 7.35 | -8.73 |

**Fig. S7.**
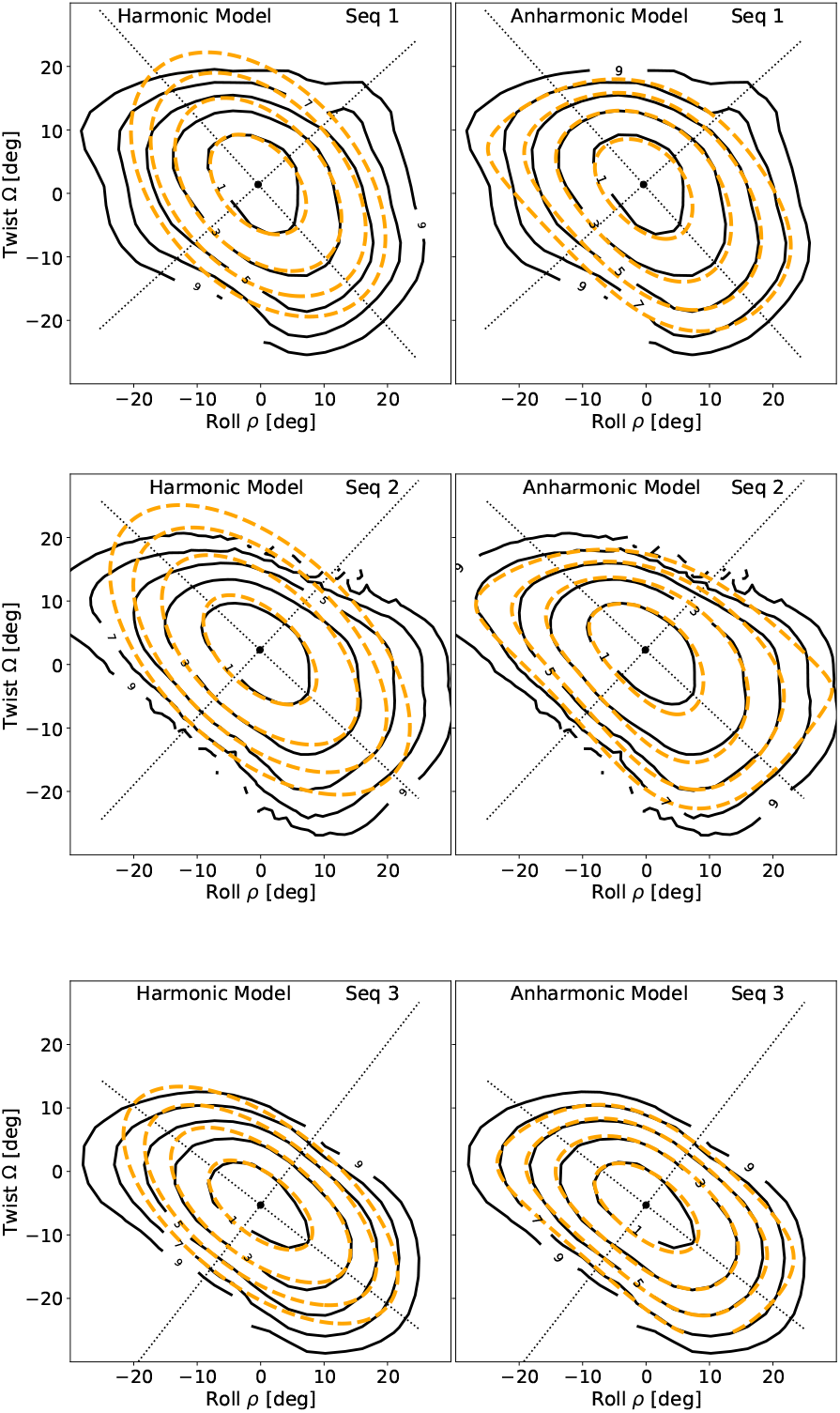
Fitted free energy landscape for sequence 1, 2, 3 (from the top to the bottom). The solid black lines are the all-atom RBB-NA data, while the dashed orange lines are the model. On the left, harmonic model (Eq. 3 of the main text, neglecting tilt) obtained fitting the all-atom data for Δ*F ≤* 3*k*_*B*_*T*. On the right side the fit to the anharmonic model (Eq. 6 of the main text) for Δ*F ≤* 7*K*_*B*_*T*. In black dashed lines the axis of the ellipses of the harmonic model.

## Notes

### Competing Interest Statement

The authors have declared no competing interest.

